# Colon-Specific Epigenetic Clocks from Minimal Features Reveal Disease-Driven Aging

**DOI:** 10.64898/2025.12.15.694318

**Authors:** Naor Sagy, Omer Bender, Daniel Z Bar

## Abstract

Epigenetic clocks estimate chronological and biological age from DNA methylation patterns, but conventional models typically train on hundreds of thousands of CpG sites and large training cohorts. We previously demonstrated that tissue-unique methylation sites change in a predictable manner upon aging and disease. Here, we demonstrate that clocks built from *tissue-unique* methylation sites enable accurate age prediction in the human colon using a compact feature set and limited training data. We trained a machine learning model on healthy colon tissue, identifying CpG sites that capture both chronological age and anatomical location (proximal vs. distal). This clock maintains high predictive performance (r = 0.978; MAE 3.9 years) while using an order of magnitude fewer sites and samples than traditional approaches. Applying the model to tissues from individuals with HIV infection, inflammatory bowel disease (IBD), and colonic polyps reveals consistent patterns of accelerated aging, while aspirin treatment is associated with partial deceleration. Our findings establish tissue-unique CpGs as a powerful basis for efficient, interpretable clocks and offer new insights into how chronic inflammation and neoplasia shape the aging landscape of the colon.

## Introduction

Aging is characterized by cumulative molecular alterations that impair tissue homeostasis and resilience ^1,2^. One of the most robust molecular signatures of aging is the progressive remodeling of DNA methylation, particularly at CpG dinucleotides ^3,4^. These age-associated methylation patterns have enabled the construction of “epigenetic clocks”: mathematical models that estimate an individual’s chronological or biological age from genome-wide methylation data. Such clocks have proven valuable in aging research and clinical studies, often correlating with age-related disease burden, frailty, and mortality risk ^5^.

Despite their utility, most widely used clocks are trained on methylation data from peripheral blood or multi-tissue datasets and rely on hundreds to thousands of CpG sites. While this broad design captures systemic aging signals, it can obscure tissue-specific biology and requires large sample sizes for accurate training. Moreover, many existing clocks are optimized for generalizability rather than precision within specific organs or disease contexts. This limits their interpretability when applied to tissues with distinct cellular compositions or exposure histories.

In contrast to these pan-tissue approaches, *tissue-unique methylation sites* represent a promising alternative for constructing efficient, biologically grounded age predictors ^6^. These are CpG sites that exhibit consistent, distinguishable methylation patterns in one tissue type compared to all others ^6,7^. They are identified through large-scale comparative analyses across multiple tissues, isolating loci with high specificity to a single tissue’s epigenetic landscape. Such sites are often enriched for enhancers of genes involved in tissue-specific function and development. Moreover, in multiple tissues these sites change in a predictable manner upon aging and disease ^6,7^. By restricting age modeling to tissue-unique CpGs, it becomes possible to train accurate predictors using far fewer features and smaller datasets, while maintaining biological interpretability.

The colon is a tissue of particular interest for epigenetic aging research. It undergoes continuous cellular renewal and is chronically exposed to environmental, microbial, and inflammatory stimuli. Epigenetic alterations in the colon are tightly linked to neoplastic transformation and inflammatory diseases such as ulcerative colitis and Crohn’s disease ^8–10^. However, existing methylation clocks provide limited insight into how aging unfolds within colon tissue specifically, or how this process may be accelerated in disease states.

In this study, we construct a colon-specific epigenetic clock using tissue-unique CpG sites. We identify colon-specific methylation features from public datasets and use these to train a machine learning model on healthy colon tissue. The resulting clock captures both chronological age and anatomical location (distal vs. proximal) with high accuracy, despite relying on a minimal set of 162 features. We then apply this model to colon and ileum samples from individuals with HIV, inflammatory bowel disease, and colonic polyps to quantify disease-associated epigenetic aging. Finally, we assess the potential mitigating effect of long-term aspirin use on methylation-based age acceleration. Our results demonstrate that clocks based on tissue-unique sites are efficient, interpretable, and sensitive to disease-driven alterations in biological aging.

## Results

### Tissue-Unique CpG Methylation Encodes Chronological Age and Colonic Region

Tissue-unique CpG sites are sites that show a distinct methylation pattern in one tissue, compared to all other examined tissues ^6,7,11^. Unique-Low (UL) sites are sites that show lower methylation values in the examined tissue, as compared to all other tissues. By contrast, Unique-High (UH) show higher methylation values than all other examined tissues. Importantly, UH and UL sites show an overlap in the range of methylation levels, and are characterized by comparison to the methylation profile of the same site in other tissues. These sites overwhelmingly regress to the mean in multiple diseases, stress and aging, a phenomenon we interpreted as epigenetic information loss ^12^. However, in some tissues, they diverge from the mean. UL sites are enriched in tissue-specific enhancers, while UH sites, at least in the liver, are enriched with causal drivers of aging ^11^. Among the CpGs included on the 27K array, 290 were classified as tissue-unique in the colon. Of these, 87% were UH and 13% were UL. Upon aging, divergence from the mean was observed in 86% of UH sites (p = 10^-40^) and 58% (p = 0.16) of UL sites. To investigate whether tissue-unique CpG sites in the colon reflect meaningful biological signals, we performed principal component analysis (PCA) on the methylation profiles of these sites across healthy control samples. The first principal component (PC1) captured a strong age-associated signal, displaying a continuous gradient across chronological age (**Fig. 1A**). In contrast, the second principal component (PC2) stratified samples by anatomical location, clearly separating proximal and distal colon tissue (**Fig. 1B**).

**Figure 1:**
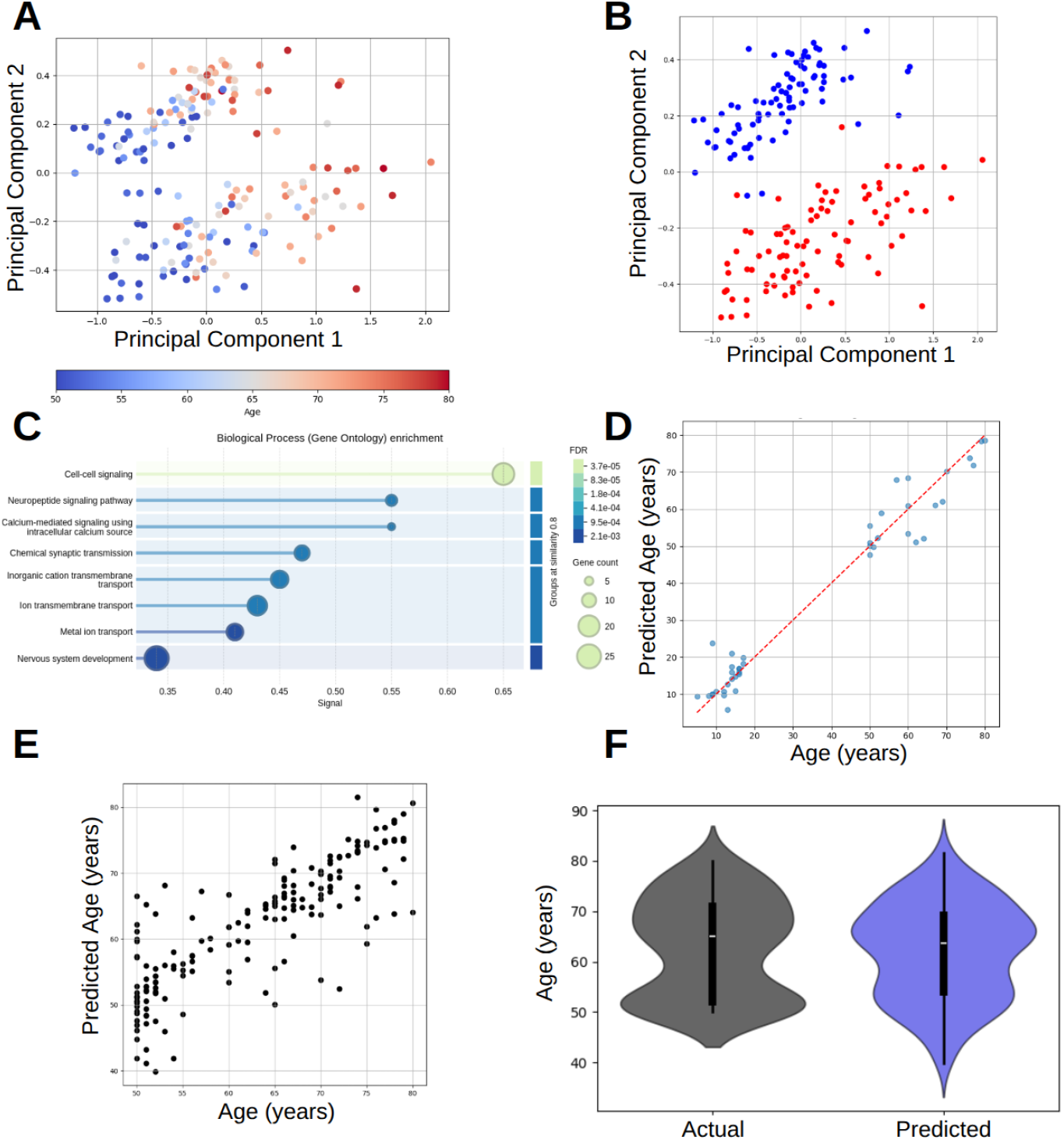
Tissue uniquely methylated sites predict chronological age. **(A)** PCA of uniquely methylated sites projected on 2D space, gradient colored by age **(B)** PC2 separates samples to distal (red) and proximal (blue) locations **(C)** STRING analysis of genes contributing to PC1 **(D)** MLP regressor model training results **(E)** Validation of MLP regressor model on an independent healthy distal colon cohort (GSE48988) **(F)** Age distribution for cohort (gray) and predicted age by MLP regressor for the same cohort (blue).

The contribution of CpG sites to both components was broadly distributed, with no single site dominating the variance (max contribution: 1.38% for PC1; 3.89% for PC2). This uniformity suggests that aging-related and spatially distinct epigenetic drift occurs across many sites in parallel. Quantification of inequality using the Gini coefficient confirmed a flatter distribution for PC1 (0.273) compared to PC2 (0.489), indicating that the aging signal is more evenly encoded across CpGs than the location-specific signature.

To better understand the functional roles of these CpG sites, we performed protein–protein interaction (PPI) enrichment analysis using STRING ^13^. Genes associated with CpGs contributing most to PC2, those separating proximal and distal regions, were enriched for cell–cell signaling pathways (PPI enrichment p = 0.0084), consistent with spatial differences in epithelial function and organization. In contrast, genes contributing to PC1, the aging axis, were enriched for a broader array of pathways including synaptic signaling, calcium-mediated communication, and cell junction integrity (PPI enrichment p = 2.2 × 10^-16^, **Fig. 1C, Supp. Table 2**). These findings suggest that the methylation drift associated with aging in the colon reflects not just random noise but functional changes in cellular signaling and barrier maintenance. Supporting this, eForge analysis of CpG sites contributing to PC1 revealed regulatory enrichment in relevant tissues composed predominantly of epithelial cell types (Mucosa layers, small intestine, sigmoid colon; **Supp. Fig. 1**), reinforcing the biological specificity of the identified CpGs ^14^.

### Tissue-Unique CpGs Enable Accurate Biological Age Prediction in Colon Tissue

Building on the observation that PC1 encodes a strong age-related signal, we trained a supervised age predictor using methylation levels at tissue-unique CpG sites from healthy colon samples. A multilayer perceptron (MLP) regressor was trained on distal colon samples pooled from multiple datasets (n = 90), using only uniquely methylated CpGs mapped to the 27K array and available in all datasets used (N = 162). Despite this compact feature space, the model achieved high accuracy on held-out test data, with a Pearson correlation of r = 0.978 (p = 5.8 × 10^-29^) and a mean absolute error (MAE) of 3.9 years (**Fig. 1D**). By contrast, applying the same model to randomly-selected sets of the same size did not produce significant learning (r=0.17, MAE = 24.7 years; **Supp. Fig. 2**)

When applied to a broader cohort of healthy distal colon samples (GSE48988), predicted biological age remained strongly correlated with chronological age (r = 0.828, p = 4 × 10^-46^, MAE = 3.8 years; **Fig. 1E**). These results demonstrate that tissue-unique CpGs, even in limited site and sample numbers, are sufficient to reconstruct individual aging trajectories within colon tissue.

### Age-Correlated Sites Significantly Overlap Tissue-Unique CpGs and Exhibit Methylation Gain

To evaluate the relationship between tissue-unique CpGs and canonical age-associated methylation drift, we identified CpG sites whose methylation levels significantly correlated with chronological age in a reference dataset of healthy colon samples (^15^; GSE48988). Of the 290 tissue-unique in the colon, 136 overlapped with the 456 age-correlated CpGs (**Supp. Fig. 3**; hypergeometric test p = 1.1 × 10^-167^), suggesting that a substantial fraction of the tissue-specific methylation landscape is also shaped by aging.

Notably, ∼99% of age-correlated sites (451 out of 456) showed a gain of methylation with age (p-value 1.6e-67; binomial test under null hypothesis *p*=0.68, as 68% of all sites are positively correlated with age; threshold p-value for a valid correlation value is p<0.05), consistent with prior observations of age-related hypermethylation. This directional bias supports a model in which functional decline in the colon is linked to epigenetic suppression at tissue-relevant loci. In contrast, CpGs that distinguish between proximal and distal colon regions maintained stable methylation patterns across age, suggesting that spatial identity is preserved independently of aging trajectories.

### HIV Infection Is Associated with Accelerated Epigenetic Aging in the Colon and Ileum

Previous studies have reported signs of accelerated biological aging in the gastrointestinal tract of individuals living with HIV, including altered methylation profiles and impaired epithelial barrier function ^16,17^. To determine whether our tissue-specific clock could detect this acceleration, we applied the model to colon and ileum samples from HIV-positive individuals on antiretroviral therapy (ART). When projecting these samples into the PCA space defined by tissue-unique CpGs, distal colon tissues from HIV-positive individuals mapped to the expected distal region along PC2, confirming anatomical consistency. However, these samples were shifted along PC1 compared to age-matched healthy controls, indicating an older biological age signature (**Fig. 2A**).

**Figure 2:**
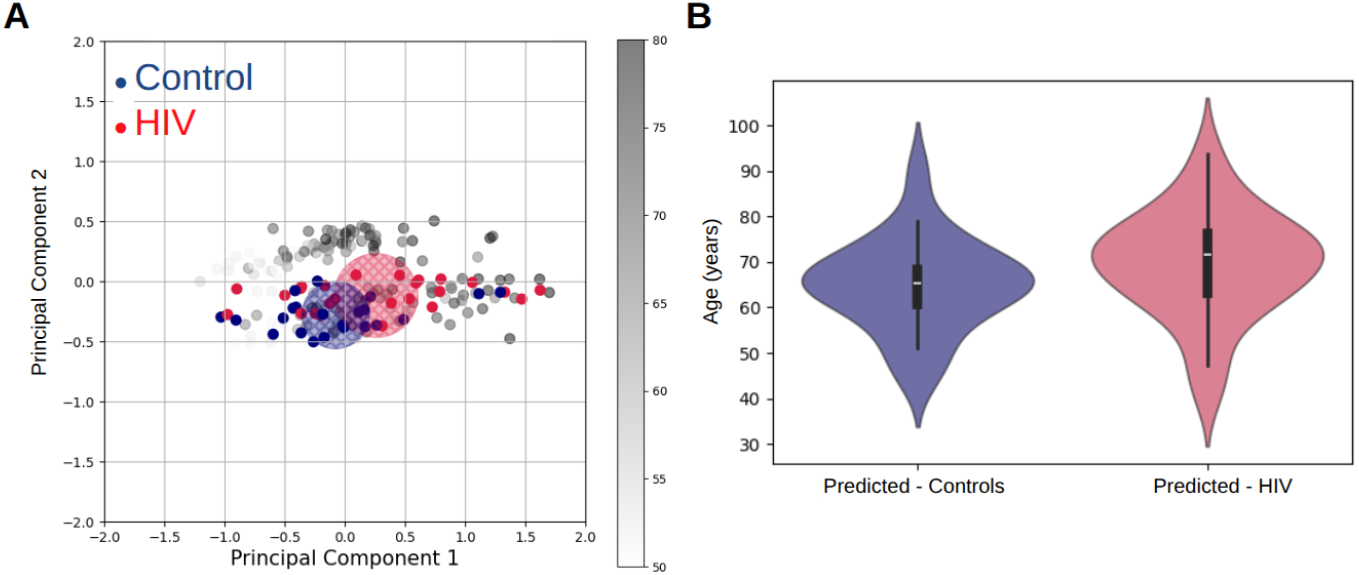
HIV accelerates age in the colon. **(A)** HIV methylation samples projected on PC1 show age acceleration **(B)** Age prediction according to methylation shows significant difference (p<0.05) between HIV positive and negative subjects.

Application of the MLP regression model confirmed this finding: colon tissue from HIV-positive individuals showed significantly higher predicted biological ages than controls (one sided Man-Whitney U test, p<0.05; **Fig. 2B**). While distal colon samples remained spatially consistent, ileum samples clustered between distal and proximal regions on PC2, consistent with their intermediate anatomical identity (**Supp. Fig. 4**). Interestingly, both colon and ileum samples from HIV-positive individuals exhibited enrichment in hypermethylation of sites associated with tight junction genes, aligning with STRING-based enrichment of cell–cell junction proteins among age-associated CpGs (**Supp. Table 2**). This suggests a possible molecular link between barrier integrity and accelerated tissue aging in the context of chronic infection.

### Ulcerative Colitis Accelerates Epigenetic Aging in Adults but Not in Pediatric Tissue

To isolate the impact of ulcerative colitis (UC) on biological aging within the colon, we analyzed DNA methylation data from monozygotic twin pairs discordant for disease. Despite their shared genetic background and identical chronological age, UC-affected twins consistently showed elevated PC1 values compared to their healthy siblings, corresponding to a mean biological age difference of 4.96 years (**Fig. 3A**; Wilcoxon signed-rank test, p = 0.032; n = 10 pairs). This result suggests that chronic inflammation in adulthood is associated with accelerated epigenetic aging of colonic tissue.

**Figure 3:**
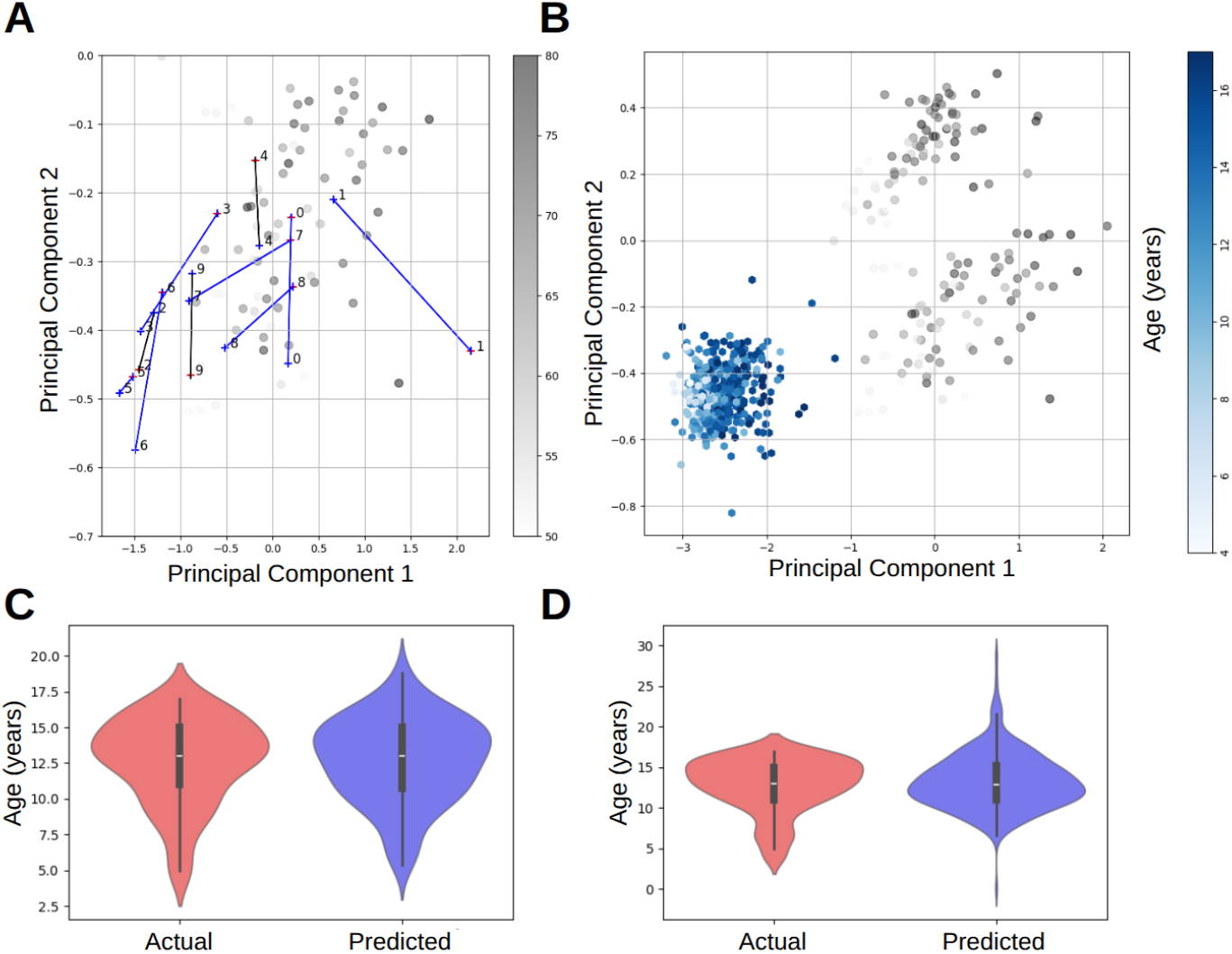
Chronic ulcerative colitis in adults, but not children, accelerates colonic epigenetic aging. **(A)** Colonic age gap in twins discordant for UC overlaid on 90 distal samples from GSE48988. Blue indicates a positive correlation of colitis and PC1. **(B)** UC-affected children (shades of blue) did not exhibit colonic age acceleration **(C**,**D)** No age acceleration revealed between predicted and real age groups for control and UC.

In contrast, when applying the same model to a large pediatric cohort (N = 296 ages 4–17) with rectal biopsies from children diagnosed with UC or non-IBD controls (^18^; GSE185061), we observed no evidence of age acceleration. The methylation profiles of UC-affected children followed the expected age gradient in PCA space (**Fig. 3B**), and predicted biological ages were statistically indistinguishable from controls (**Fig. 3C,D**). These findings indicate that age acceleration is not a universal consequence of colitis, but may emerge with disease chronicity or in the context of age-related epigenetic plasticity in adult tissues.

### Colonic Polyps Exhibit Profound Epigenetic Age Acceleration

To evaluate whether premalignant lesions also exhibit features of accelerated aging, we applied the colon-specific clock to methylation data from colonic polyps. Despite being histologically benign, polyp samples showed marked epigenetic age acceleration, with predicted ages averaging 33.2 years older than chronological age (t-test p = 7 × 10^-26^; **Fig. 4A,B**). When projected into PCA space, polyp samples align with the canonical aging trajectory defined by healthy colon tissue, suggesting that they follow the same epigenetic program but are shifted forward along the age axis.

**Figure 4:**
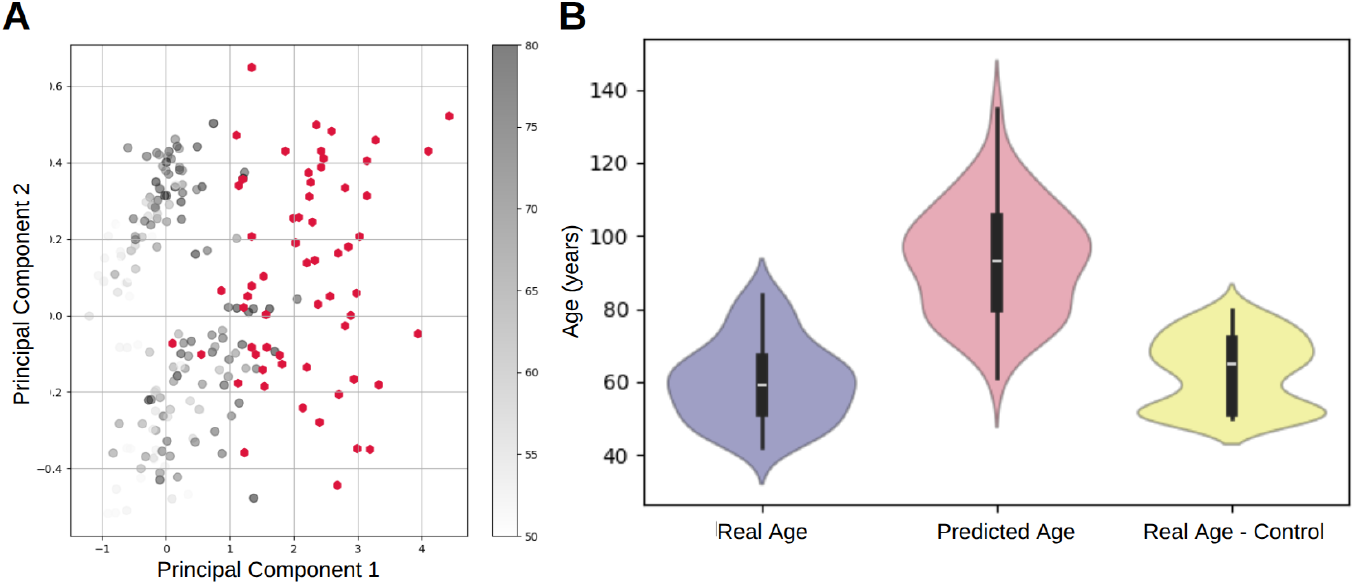
Colonic polyps follow canonical aging trajectory, but are epigenetically older. **(A)** Mapping polyps methylation into PCA 2D space **(B)** Age violin plots for real age, predicted age, and the control group with which the PCA transformation was computed.

Interestingly, the degree of age acceleration did not correlate with reported histological grade or subtype, implying that the observed methylation aging signal reflects a generalized molecular shift rather than lesion-specific changes (**Supp. Fig. 5A,B**). These results suggest that polyp formation may be accompanied, or even preceded, by extensive epigenetic aging, potentially priming the tissue for neoplastic transformation.

### Long-Term Aspirin Use Is Associated with Reduced Epigenetic Aging in the Colon

Aspirin is widely recognized for its anti-inflammatory effects and has been associated with reduced risk of colorectal cancer in both epidemiological and interventional studies ^19,20^. Its protective role is thought to involve modulation of prostaglandin signaling, immune activity, and epithelial turnover in the gut. To assess whether long-term aspirin use also influences molecular aging, we applied our colon-specific epigenetic clock to distal colon samples from individuals with and without chronic aspirin exposure.

Predicted biological age was significantly lower in aspirin users compared to age-matched non-users (Mann–Whitney U test, p = 0.033; n_1_ = 11, n_2_ = 20), suggesting that long-term aspirin treatment may decelerate age-associated methylation changes in colonic tissue (**Fig. 5**). This shift occurred despite similar chronological age distributions across groups, indicating a potential protective effect of aspirin against molecular aging in the colon. While causality cannot be inferred from these cross-sectional data, the finding aligns with aspirin’s broader role in preserving epithelial integrity and suppressing inflammation-driven damage over time.

**Figure 5:**
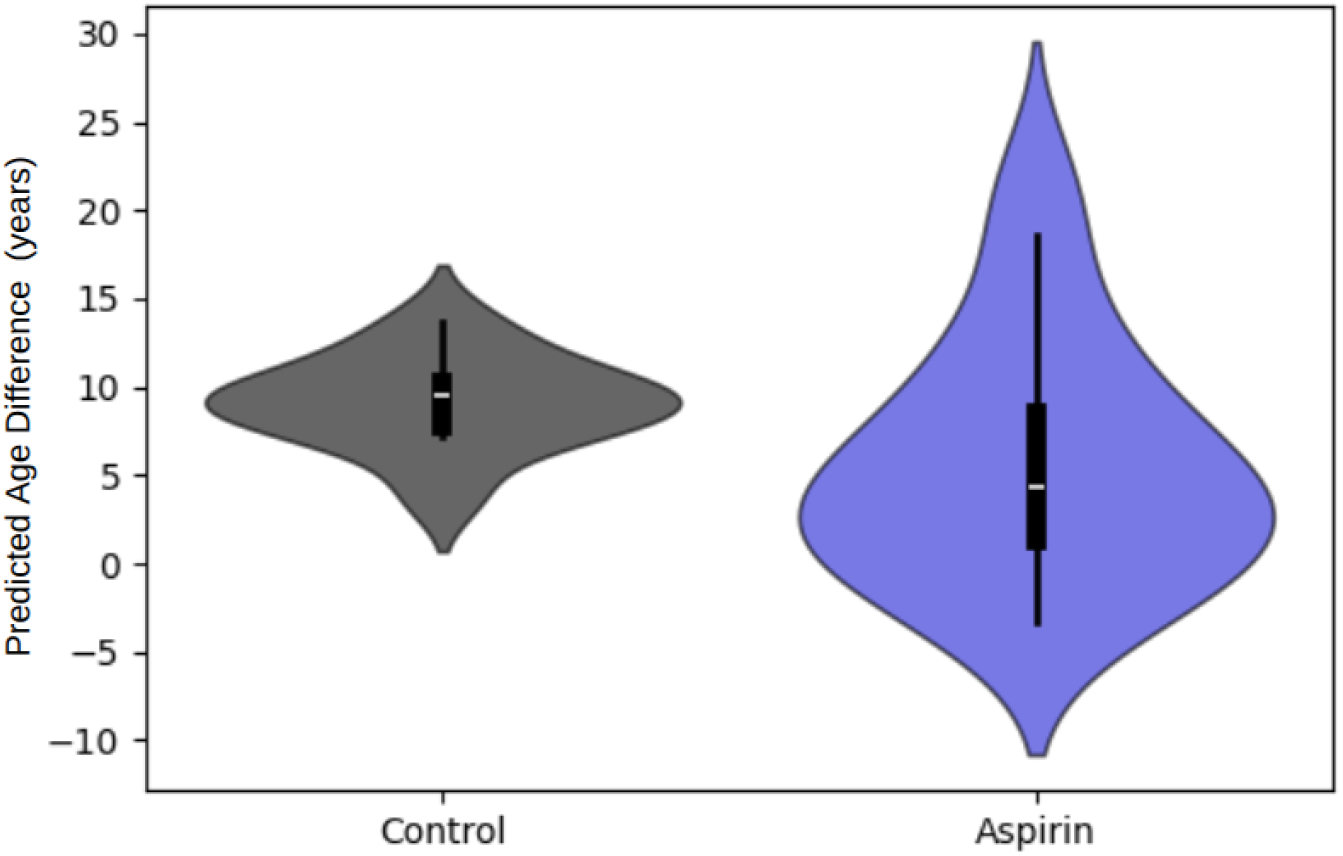
Reduced colon epigenetic age in long-term aspirin users. Difference between actual and predicted age in non-treated (Control) and aspirin treated (Aspirin) individuals. * - p < 0.05.

### FAIRE colon-unique sites reveals

To gain mechanistic insight into the nature of these tissue-unique marks, we analyzed DNA accessibility in colon tissue using the Formaldehyde-Assisted Isolation of Regulatory Elements (FAIRE; ^21^) technique. The analysis revealed a stark contrast between UL and UH sites. FAIRE density signals peaked sharply around UL sites, an indicator of highly accessible, open chromatin. In dramatic opposition, the FAIRE signal dropped significantly at UH sites, suggesting a closed chromatin structure. Importantly, this difference in chromatin accessibility was not merely a consequence of the sites’ methylation levels (UL average 0.62 vs. UH average 0.4 beta values; Welch’s t-test, p = 1.3 × 10^-265^), where the difference is in the opposite direction of classical methylation-accessibility models. This structural finding aligns perfectly with our functional results, as UL sites in digestive tract tissues were significantly enriched for active enhancers (**Fig. 6C, Supp. Fig. 6**), similarly to what we previously shown in liver and kidney ^6^, establishing a clear link between tissue-unique low methylation and a biologically critical, open chromatin state.

**Figure 6:**
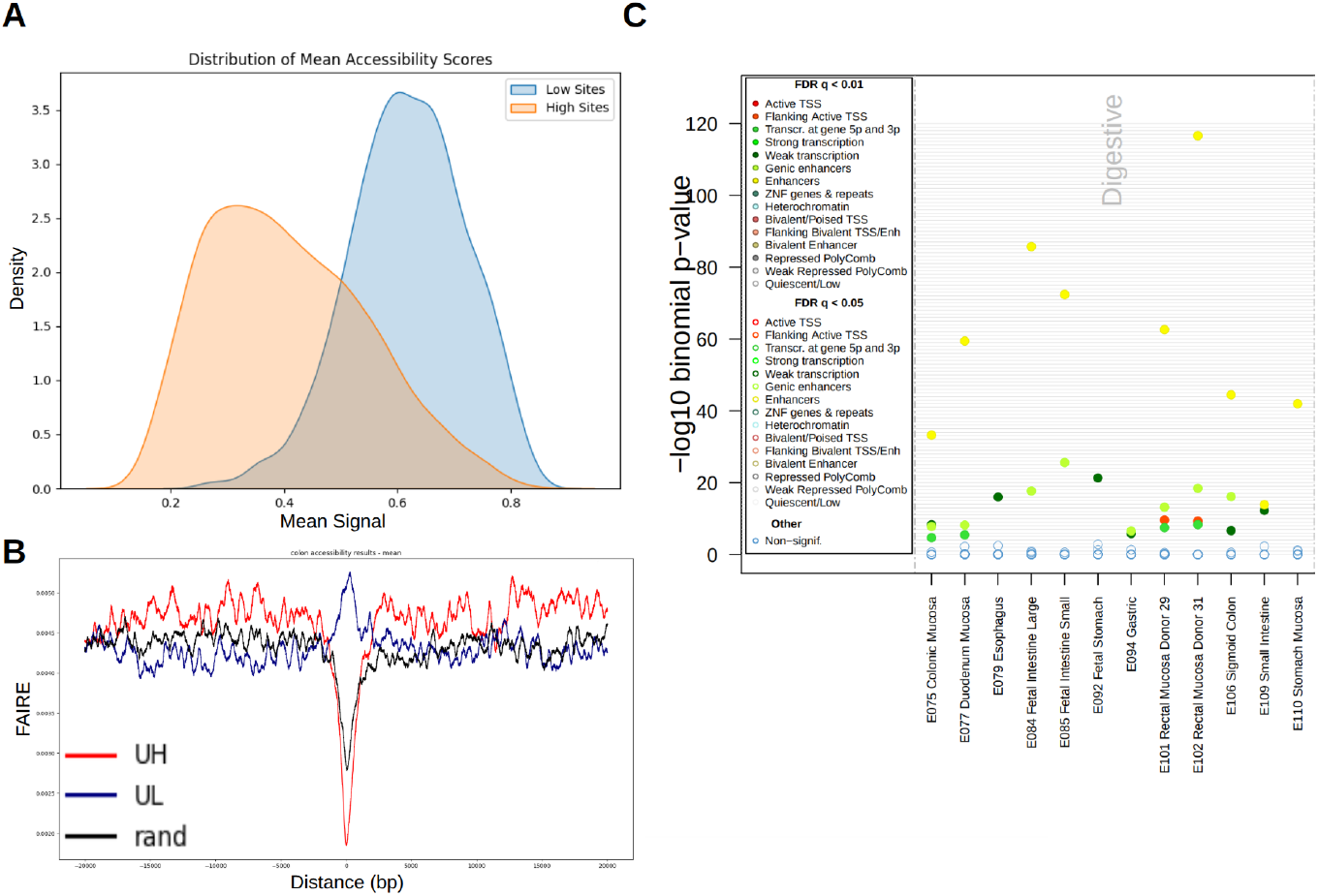
FAIRE analysis reveals distinct chromatin accessibility patterns at UL and UH CpG sites in colon tissue: **(A)** UH and UL sites methylation density distribution in colon **(B)** FAIRE signal analysis for UH and UL **(C)** eFORGE enrichment analysis for UL sites, showing enhancer (yellow circles) enrichment in the digestive tract. Cells of other origins did not show this phenomenon, indicating tissue-specific enhancers.

## Discussion

In this study, we demonstrate that tissue-unique CpG sites in the human colon encode sufficient information to construct accurate, biologically meaningful epigenetic clocks. By focusing exclusively on CpGs with tissue-specific methylation patterns, we were able to train a high-performing age predictor using substantially fewer features and smaller sample sizes than conventional models. This compact, interpretable framework enabled us to uncover distinct aging trajectories in healthy colon tissue and to detect significant age acceleration in multiple disease contexts, including HIV infection, ulcerative colitis, and colonic polyps.

Our approach differs from previous clocks in both design and intent. Most epigenetic clocks rely on agnostic feature selection across the entire methylome, often prioritizing predictive power over biological specificity. By contrast, we anchor our model in tissue-unique CpGs, sites defined by their specificity to the colonic epigenome relative to other tissues. Mechanistically, these sites are biologically coherent: Functional analysis of DNA accessibility (FAIRE) showed that Unique-Low (UL) sites are highly accessible (peaked FAIRE signal), consistent with enrichment in tissue-specific enhancers, while Unique-High (UH) sites show a pronounced drop in accessibility. Moreover, the vast majority of age-correlated sites in our analysis exhibited a gain of methylation (hypermethylation) with age, suggesting a molecular link between epigenetic suppression at these tissue-relevant loci and age-related functional decline. This grounding in specific chromatin states and directional change makes our minimal-feature clock highly interpretable.

The aging signal captured by our model is not only statistically robust but also biologically coherent. Principal component analysis revealed that tissue-unique CpGs stratify samples by both chronological age and anatomical location (distal vs. proximal), reflecting the structured, spatially encoded nature of epigenetic drift in the colon. Importantly, disease-associated samples did not deviate from this trajectory but instead appeared “older” along the same axis, suggesting that chronic inflammation, infection, and neoplastic change accelerate rather than reprogram the aging process.

The application of our clock provided quantitative estimates of disease-driven age acceleration. Colonic polyps exhibited the most profound shift, with a predicted epigenetic age averaging 33.2 years older than the chronological age. This dramatic acceleration suggests that pre-malignant lesions are accompanied by extensive epigenetic aging, potentially creating a permissive landscape for neoplastic transformation. Similarly, the chronic inflammation of Ulcerative Colitis (UC) accelerated epigenetic aging in adult discordant twin pairs (mean difference of 4.96 years), yet no such acceleration was found in pediatric UC patients. This crucial contrast suggests that age acceleration is not a universal consequence of colitis but may require age-related epigenetic plasticity or disease chronicity to emerge in adult tissue.

Finally, we explored a potential mitigating factor against this molecular aging: long-term aspirin use. Our data revealed that predicted biological age was significantly lower in chronic aspirin users compared to age-matched non-users. This finding aligns with aspirin’s known anti-inflammatory and cancer-protective roles in the colorectum and suggests a partial deceleration of age-associated methylation changes in the colonic tissue. While causality requires further investigation, this result points to a clear protective epigenetic signature associated with the drug.

## Methods

### Identification of human colon-unique sites

Colon-unique methylation sites were taken from ^7^. A unique high methylation site in the colon is defined as a site where the average methylation value is higher by 0.1 or more than the 0.95 quantile of all other tissues, and a unique low methylation site is one where the average methylation value is lower by 0.1 or more than the 0.05 quantile of all other tissues.

### Datasets used

Methylation data were obtained from the Gene Expression Omnibus. All datasets used are listed in **Supplementary Table 1**. Analysis was conducted on the processed Series Matrix Files, which were separated into a methylation data file and an experiment metadata files.

### Principal component analysis

PCA linear dimensionality reduction was employed to project n-dimensional data (n CpG sites) onto 2D space. We utilized the Python scikit-learn package to perform PCA. The data were parsed and rearranged from a dataframe format to comply with the package’s format requirements, representing probes as dimensions. Whenever a subset of the database was required, we either reconstructed the input files to include only the required dataset or filtered out test subjects in the code prior to implementing PCA on the data. Filtering was conducted on metadata stored in a separate dataframe and/or on methylation sites. Projecting samples on baseline PCA space to show age drift was performed by transforming test samples with the already fitted PCA model on normal colon samples.

### Model training

We employed a multilayer perceptron (MLP) regressor using Python package scikit-learn 1.6.1 to model the data; The network architecture consisted of two hidden layers with 100 and 50 neurons respectively. The ReLu activation function was used and the model was optimized with Adam solver. Training was performed for a maximum of 1000 iterations; The dataset was split into training and test sets using a 80%-20% ratio.

### Age correlation and unique sites overlap

Correlation between each CpG site methylation and age was calculated per site on GSE48988, dropping CpG sites with less than 25 valid methylation data points. A hypergeometric statistical test was applied to quantify unique sites and age correlated sites overlap significance. To accommodate chip size only unique sites mapped to the 27K chip used in GSE48988, yielding 290/3893 unique sites.

### DNA accessibility analysis

To compare DNA accessibility between different types of CpG sites, we downloaded the FAIRE density signal for colon tissue from the UCSC Genome Browser (accession: wgEncodeEH003495). We randomly selected 500 CpG sites from each of three categories: uniquely high, uniquely low, and random. For each site, a 16,384 bp region centered on the CpG was extracted from the FAIRE signal track. The accessibility scores across these regions were averaged within each group to assess overall patterns of chromatin accessibility. The above analysis was performed using Python 3.12.

### Methylation enrichment analysis

eFORGE 2.0 (Experimentally derived Functional element Overlap analysis of ReGions from EWAS; ^22^, https://eforge.altiusinstitute.org/) was used to assess the functional and tissue-specific relevance of differentially methylated colon-unique CpG sites. Analyses were performed with a 1 kb window around each CpG and evaluated against the 15-state ChromHMM chromatin model from the Roadmap Epigenomics and ENCODE datasets. Separate runs were conducted for UH and UL sites. Enrichment significance was assessed using the eFORGE 2.0 statistical framework, with a strict p-value threshold of 0.01 for high-confidence hits and a marginal threshold of 0.05 for suggestive associations.

## Data and materials availability

All methylation data used came from previously published sources, identified and references in the text and supplementary files. Generated data are available as supplementary files.

## Acknowledgments

We thank the Bar lab members for comments and suggestions. This work was supported by the Israeli Science Foundation (grants 654/20 and 632/20 to DZB).

## Author contributions

DZB and NS designed the experiments. NS performed the analysis. DZB and NS wrote the manuscript with critical inputs and comments from OB.

## Supplementary figure legend

**Supplementary Figure 1**

**eForge analysis of CpG sites contributing to PC1**. High resolution file attached.

**Supplementary Figure 2.**
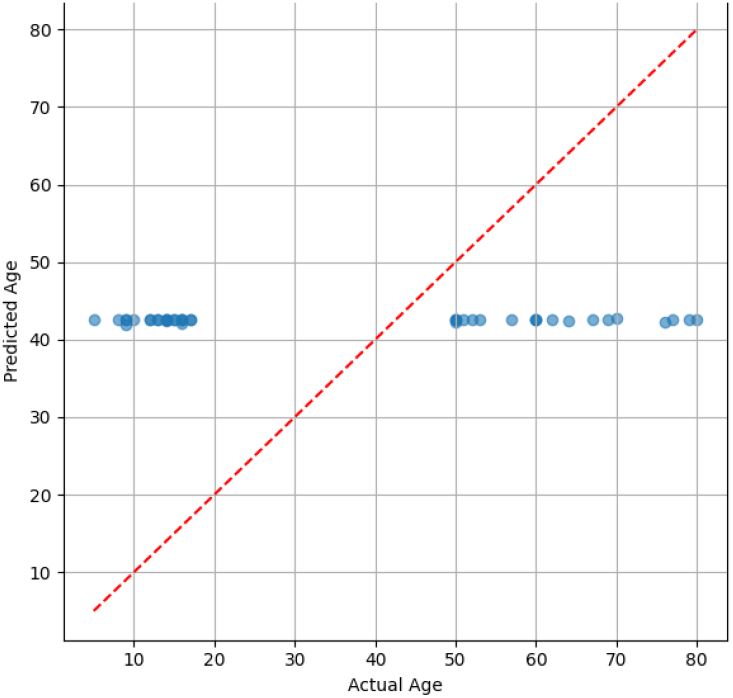
Training the MLP Regressor Model on Randomly Selected CpG Sites Fails to Predict Age. The multilayer perceptron (MLP) regressor model, when trained on a randomly selected subset of methylation sites (same size as the tissue-unique set), shows no significant correlation with chronological age.

**Supplementary Figure 3.**
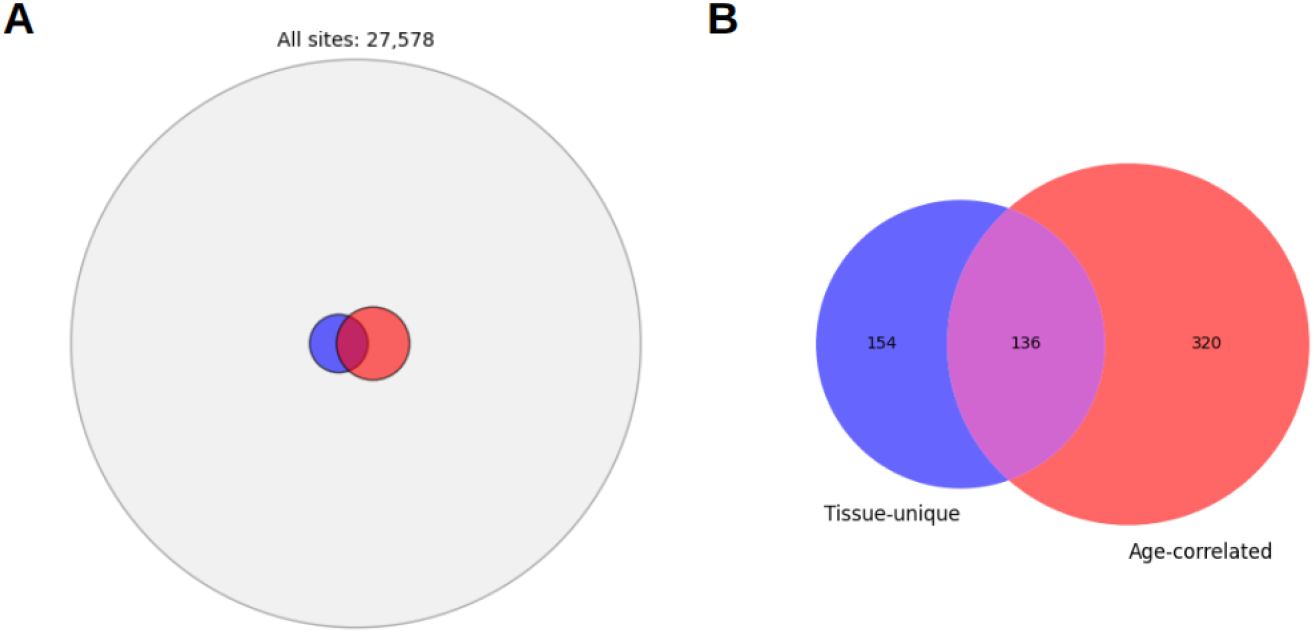
Significant Overlap Between Colon-Unique and Age-Correlated CpG Sites. **(A)** Venn diagram illustrating the overlap between the 290 tissue-unique CpG sites in the colon (blue) and the 456 CpG sites whose methylation levels significantly correlate with chronological age (Age-Correlated CpGs) within the relevant 27,578 CpG sites on the 27K array. **(B)** Zoom-in view focusing on the tissue-unique and age-correlated CpG sites.

**Supplementary Figure 4.**
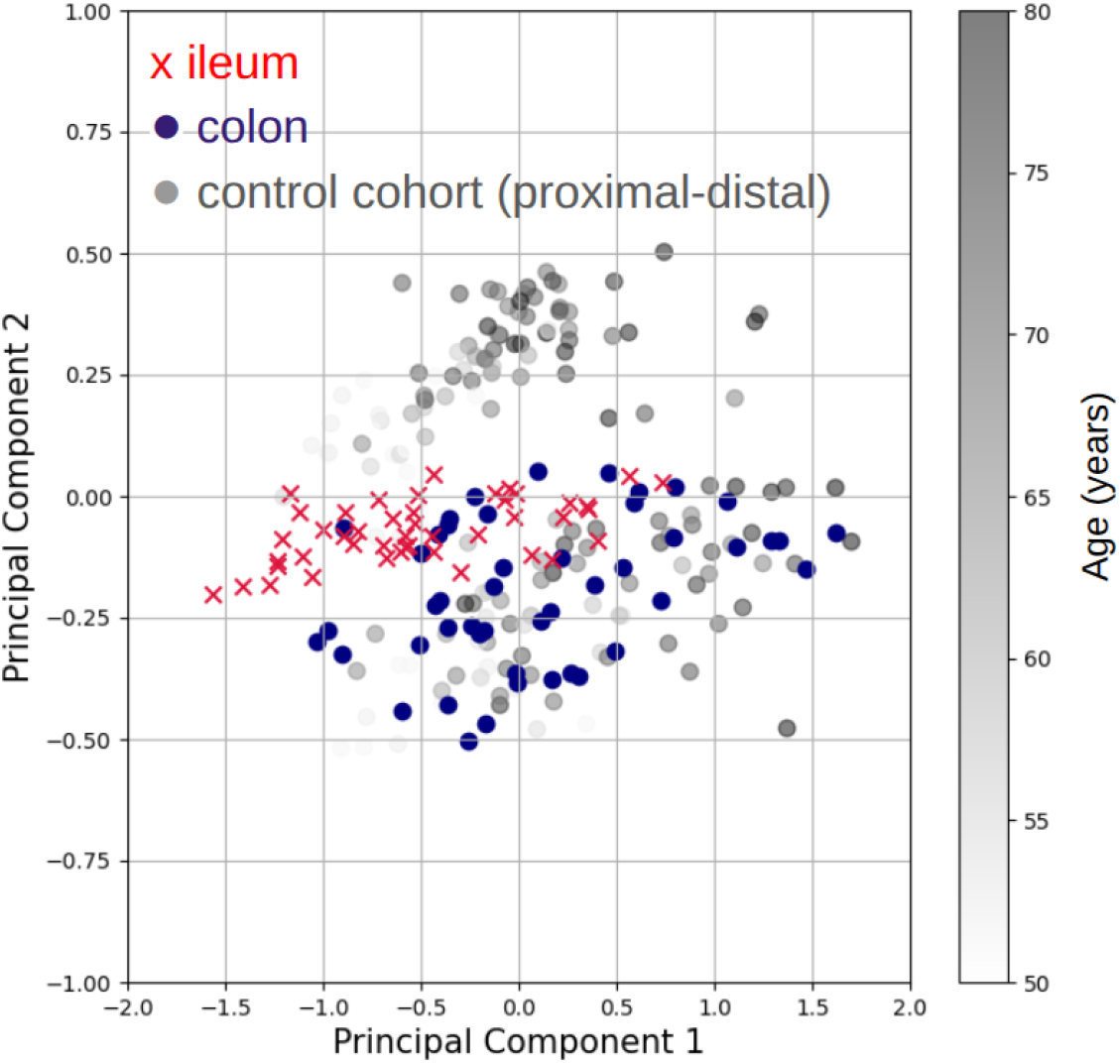
PCA Projection of Samples Confirms Anatomical Consistency of Ileum Tissue. Principal Component Analysis (PCA) projection of colon (blue) and ileum (red) samples onto the space defined by tissue-unique CpGs in healthy colon tissue. While colon samples map to the distal colon region, ileum samples cluster between the proximal and distal regions along the PC2 axis, confirming their expected intermediate anatomical identity.

**Supplementary Figure 5.**
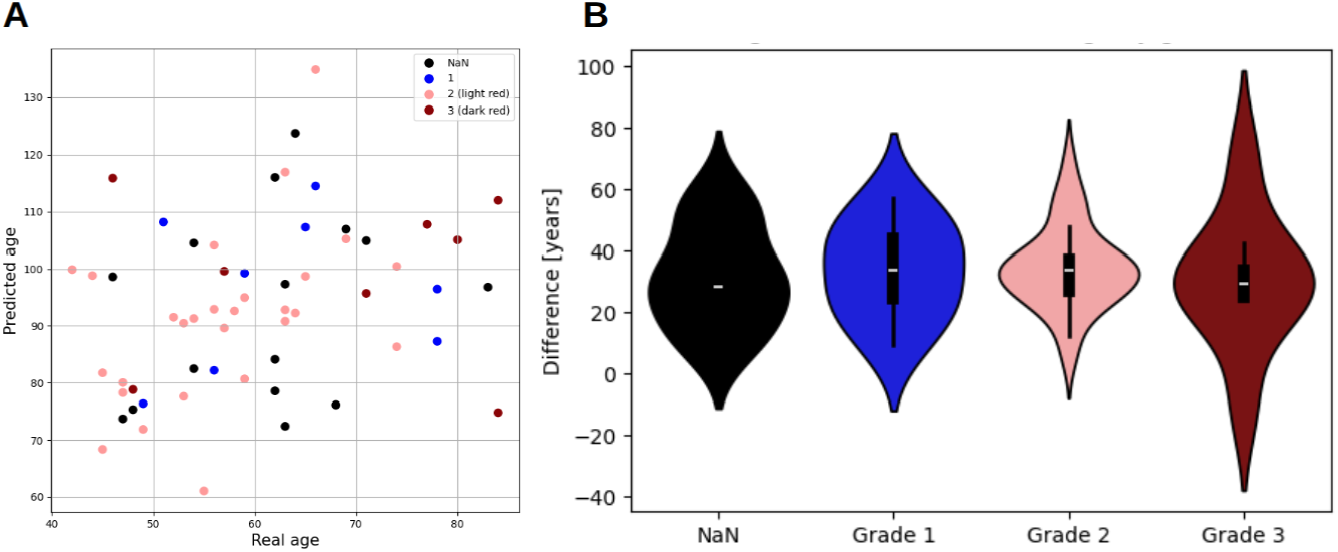
Lack of Correlation Between Epigenetic Age Acceleration and Colonic Polyp Subtype. PCA **(A)** and MLP difference from real age **(B)** analysis showing that the degree of epigenetic age acceleration observed in colonic polyp samples does not significantly correlate with the reported histological grade or subtype of the lesion. NaN - grading score adenoma not available.

**Supplementary Figure 6**

**Full eForge analysis of colon UL sites, including data not shown in Fig. 6C**. High resolution file attached.

**Supplementary Table 1.** Datasets used.

| Database | n | age range<br>[years] | mean<br>age[years] | female-male<br>ratio [%] |
| --- | --- | --- | --- | --- |
| GSE48988 | 178 (90 distal, 88 proximal) | 50-80 | 62.8 | 100-0 |
| GSE185061 | 296 (211 UC, 85 non-IBD) | 4-17 | 12.8 | 51-49 |
| GSE142257 | 62 (31 distal, 31 proximal) | 50-80 | 62.5 | 100-0 |
| GSE42921 | 23 (12 normal, 6 UC, 5 Crohn's) | 5-19 | 13.6 | 61-39 |
| GSE233603 | 61 | 42-84 | 60.3 | 48-52 |
| GSE245924 | 129 (47 colon, 44 ileum, 38 PBMC) | no age data | no age data | 20-80 |
| GSE27899 | 20 (10 UC, 10 normal) | no age data | no age data | no gender data |

**Supplementary Table 2.** PC1 PPI enrichment.

|  | function/component | enrichment p-value |
| --- | --- | --- |
| biological process | cell-cell signaling | 0.00037 |
|  | neuropeptide signaling pathway | 0.0101 |
|  | calcium-mediated signaling | 0.0139 |
|  | chemical synaptic transmission | 0.0124 |
|  | inorganic cation transmembrane transport | 0.0115 |
| cellular component | integral component of plasma membrane | 0.00042 |
|  | synapse | 0.0089 |
|  | cell junction | 0.0125 |
|  | cell periphery | 0.0089 |

