## Supplementary figures and images for "Colon-Specific Epigenetic Clocks from Minimal Features Reveal Disease-Driven Aging"

### Supp. Fig. 1

#### DMPs analyzed across samples for erc2−chromatin15state−all Unnamed

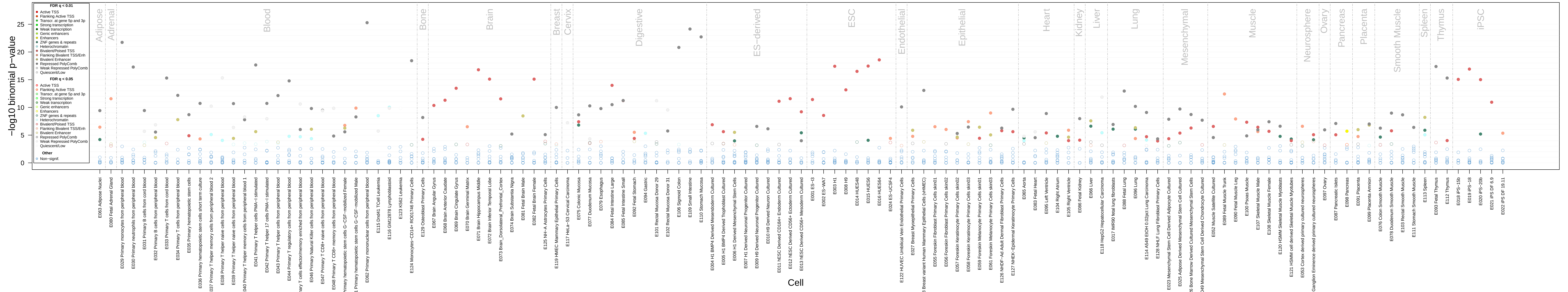

### Supp. Fig. 6

#### DMPs analyzed across samples for erc2−chromatin15state−all Unnamed

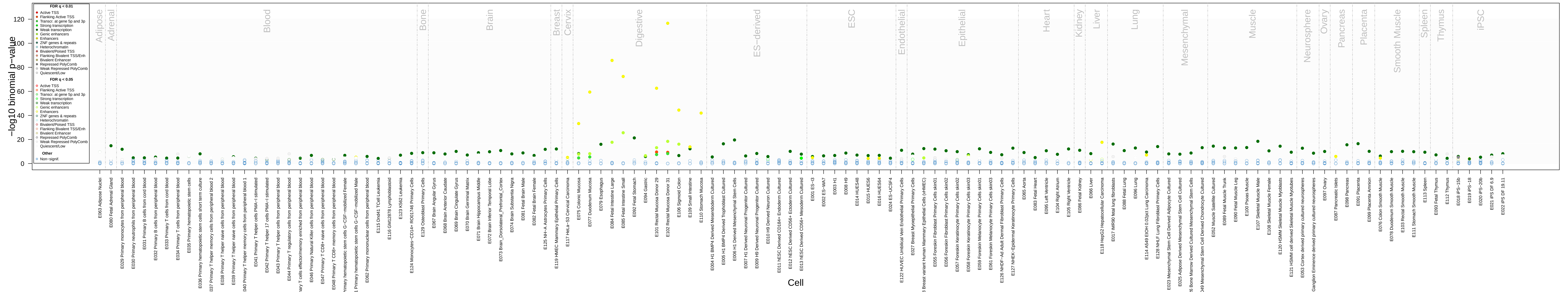
